# Evolutionary rescue is promoted in compact cellular populations

**DOI:** 10.1101/2022.05.27.493727

**Authors:** Serhii Aif, Nico Appold, Lucas Kampman, Oskar Hallatschek, Jona Kayser

## Abstract

Mutation-mediated drug resistance is one of the primary causes for the failure of modern antibiotic or chemotherapeutic treatment. Yet, in the absence of treatment many drug resistance mutations are associated with a fitness cost and therefore subject to purifying selection. While, in principle, resistant subclones can escape purifying selection via subsequent compensatory mutations, current models predict such evolutionary rescue events to be exceedingly unlikely. Here, we show that the probability of evolutionary rescue, and the resulting long-term persistence of drug resistant subclones, is dramatically increased in dense microbial populations via an inflation-selection balance that stabilizes the less-fit intermediate state. Tracking the entire evolutionary trajectory of fluorescence-augmented “synthetic mutations” in expanding yeast colonies, we trace the origin of this balance to the opposing forces of radial population growth and a clone-width-dependent weakening of selection pressures, inherent to crowded populations. Additionally conducting agent-based simulations of tumor growth, we corroborate the fundamental nature of the observed effects and demonstrate the potential impact on drug resistance evolution in cancer. The described phenomena should be considered when predicting the evolutionary dynamics of any sufficiently dense cellular populations, including pathogenic microbial biofilms and solid tumors, and their response to therapeutic interventions. Our experimental approach could be extended to systematically study rates of specific evolutionary trajectories, giving quantitative access to the evolution of complex adaptations.

---

Many drug resistance mutations are associated with a fitness cost in the absence of treatment and thus subject to purifying selection [1–4]. In principle, slower growing resistant clones can be rescued by acquiring subsequent compensatory mutations, off-setting their resistance-associated fitness cost [2, 5– 8](Fig. 1a). However, the short lifetime and small size of less-fit intermediate subclones makes crossing such a “fitness valley” inherently rare, requiring large populations [9, 10], environmental shifts [11] or long times [12] to manifest at all. We and others have recently demonstrated that in dense populations collective cell dynamics inherently decrease the power of selection by several orders of magnitude [13, 14]. The ensuing alterations in evolutionary dynamics and clone longevity are likely to also affect evolutionary rescue. However, tracking the ongoing evolutionary trajectories of small individual subclones is extremely difficult. Time-resolved deep sequencing approaches can identify emerging *de novo* sub-clones but lack the ability to spatio-temporally track small subclones, especially in dense populations [15– 17]. Microscopy-based clonal tracking approaches of fluorescently pre-labeled subclones, while featuring exquisite spatio-temporal resolution, fail to capture newly arising mutant clones [13, 14, 18–20]. Consequently, we still lack empirical insight into how the altered evolutionary dynamics in dense populations impacts subclonal evolutionary rescue and, as a result, drug resistance evolution. A better understanding of the fundamental processes governing evolutionary rescue in such crowded settings will be crucial to understand why drug resistance is so prevalent in many pathological cellular populations. In this work, we study the evolutionary rescue of drug resistant subclones in densely-packed cellular populations via a genetically tailored yeast-based model systems. Combining our empirical results with accompanying *in silico* models of colony growth and agent-based tumor simulations, we find that the competing effects of mechanically-driven radial range expansion and a clone-width dependent reduction of selection pressures conspire to create a previously unidentified inflation-selection balance. The resulting stabilization of less-fit resistant clones drastically enhances their chance to acquire compensatory mutations, persist for long times and, upon drug application, become the seed for resurgent growth.

**Figure 1:**
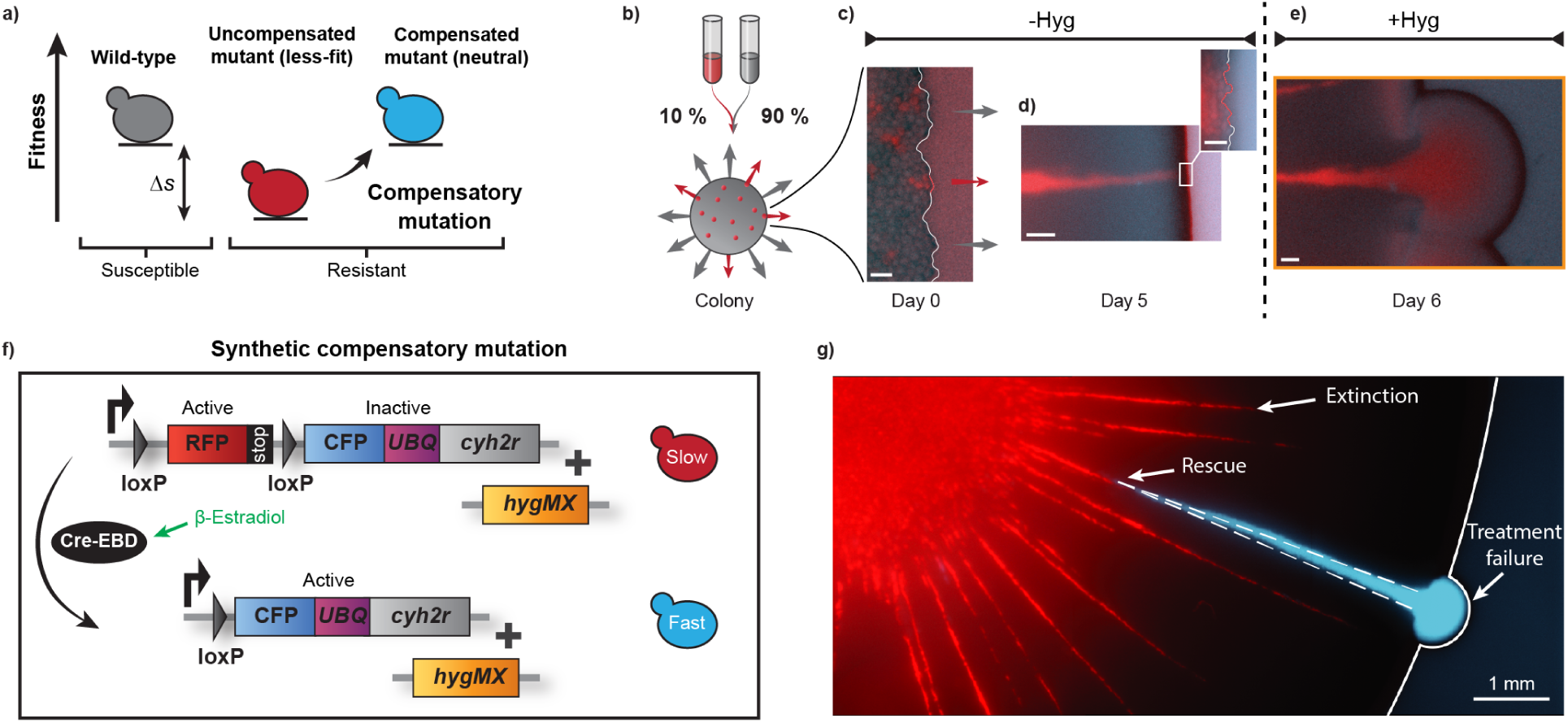
Tracking evolutionary rescue dynamics via synthetic compensatory mutations. **a)** Schematic of fitness valley crossing via compensatory mutation. The y-axis represents the fitness of the mutants, x-axis - susceptibility to the treatment drug. Grey “wild-type” strain is susceptible to the treatment, red “uncompensated mutant” has lower fitness compared to the “wild-type” and is resistant, and blue “compensated mutant” - has the same fitness as the “wild-type” and is resistant. **b)** Schematic representation of experimental competition assay: initial colonies are produced by inoculating 1 L of the mixture of 10 % “uncompensated mutant” (red) and 90 % “wild-type” (grey) onto agar plates. Microscope images of the colony front at different times of colony growth, ≈ 1 hr after inoculation (day 0) **c)**, before treatment application after 5 days of growth (day 5) **d)**, and after treatment (day 6) **e)**. Red fluorescent is “uncompensated” less-fit mutant, dark - “wild-type”. Scale bars for single-cell resolution images are 10 microns, for day 5 and 6 - 100 microns. **f)** Mutants genomes represent synthetic mutation via Cre-recombinase. (See Supplementary Video 1 for single-cell mutation event). **g)** Image of part of the colony one day after treatment application. Red clones are outcompeted by the wild-type, a mutated clone persists and leads to a “treatment failure” event, and the surrounding wild-type is halted.

## Results

### Tracking evolutionary rescue dynamics via synthetic compensatory mutations

In this study, we investigate the evolutionary rescue of slower-growing resistant subclones, growing in a background of fitter but susceptible wild-type cells. In this context, evolutionary rescue refers to the *de novo* acquisition of a compensatory mutation that elevates the fitness of resistant cells to that of the wild-type (Fig. 1a). Using a model system of multi-type yeast colonies, comprised of genetically tailored resistant and susceptible *S. cerivisiae* strains, allows us to track evolutionary rescue dynamics and the resulting fate of resistant lineages with subclonal resolution. Fluorescently labeled resistant cells, carrying a constitutive resistance against the drug Hygromycin B, are interspersed at a low fraction (10%) into a population of Hygromycin-susceptible wild-type cells (Fig. 1b). We then grow radially expanding colonies from a small droplet of this mixed inoculum, placed on a 2D agar substrate (Fig. 1c). Resistant lineages form well-segregated sectors that can be readily visualized via time-resolved fluorescence microscopy (Fig. 1d). Subjecting the population to Hygromycin after several days of Hygromycin-free growth, arrests the growth of susceptible wild-type cells (see Supplementary Fig. S1). However, persisting resistant clones seed rapidly expanding, resurgent growth cones that are unresponsive to continued treatment (Fig. 1e).

To emulate the cost of resistance, we selectively adjust the growth rates of resistant cells via the translational inhibitor cycloheximide, to which wild-type cells are insensitive.(Fig. 1f,g) [21]. Note that in the context of this study, cycloheximide is solely used to adjust the fitness cost of resistant cells, while the resistance itself refers to Hygromycin B. As a result of this fitness cost, resistant subclones remain small and are gradually expelled from the expanding front during an initial Hygromycin-free growth phase(Fig. 1d). Upon application of Hygromycin, halting the growth of susceptible wild-type cells, persisting resistant clones seed rapidly expanding, resurgent growth cones that are unresponsive to continued treatment (Fig. 1e).

To additionally detect compensatory mutation events and track the evolutionary trajectories of emerging *de novo* compensated subclones, we introduce a recombinase-based “synthetic mutations” system (Fig. 1f). Slower-growing resistant cells can stochastically switch at a rate *μ* from a slower-growing, red-fluorescent initial state to a cyan-fluorescent “rescued” state which has a growth matching that of the non-switching wild-type cells. The mutation rate *μ* can be tuned via β-estradiol induction while the emulated fitness cost of resistance Δ*s* is set by the concentrations of β-estradiol and cycloheximide, respectively (See Supplementary Fig. S2). Note that both uncompensated and compensated cells carry the Hygromycin resistance.

These synthetic mutations allow us to monitor incoming compensatory mutations throughout colony expansion and directly measure their effects on long-term persistence and treatment failure. With each initial clone representing an independent experiment, this high-throughput approach allowed us to quantitatively assess the fate of more than *N* = 10000 resistant lineages in parallel. Fig. 1g shows a representative post-experiment image of a colony with clone boundaries as a “fossil record” of past clone widths. We find that slower-growing resistant clones form narrow yet surprisingly persistent streaks before eventually being expelled from the expanding front. A compensated resistant subclone, in contrast, can persist long-term, even grow in size and eventually seed a resurgent growth cone upon Hygromycin treatment.

### Dynamics of clone width distribution and rescue efficacy

To quantify the underlying evolutionary dynamics, we recorded the complete history of *N* ≳ 3000 individual resistant subclones - initially associated with an intrinsic fitness cost of *s* = 0.013 *±* 0.006 and a compensatory mutation rate of 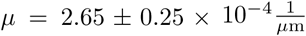 - via time-resolved, multi-scale fluorescence microscopy in 24h intervals (Fig. 2a). Segmenting images via a machine-learning-based, pixel-wise segmentation pipeline, we measured the width and compensation state (uncompensated (-) or compensated (+)) of clones at the colony edge to reconstruct their complete evolutionary trajectory (see Fig. 2b).

**Figure 2:**
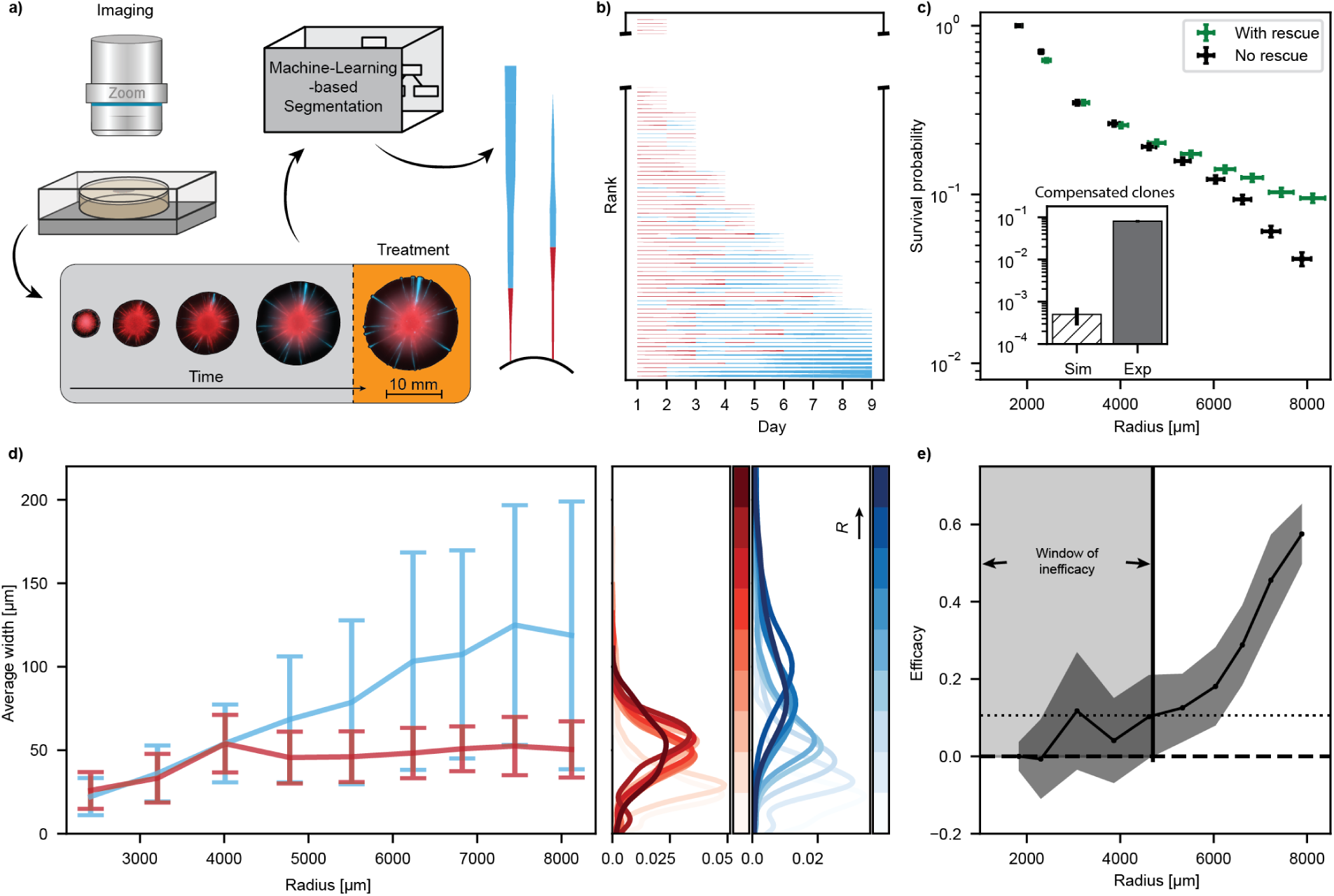
Dynamics of clone width distribution and rescue efficacy in experiments. **a)** Experimental data analysis pipeline. Time series of colony images pre and post-treatment obtained by zoom microscope and processed via machine-learning-based image segmentation to extract clone trajectories. **b)** Illustration of full clone’s history from 1 to 9 days of growth of one colony. Line width is proportional to the clone width of the lineage at the colony’s edge. The broken axis represents that more uncompensated clones were extinct at day 2 for this colony, but are not shown (see Supplementary Fig. S3 for trajectories of no rescue control). **c)** Clone survival probabilities at the front (Eq. 1). Green markers - experiments with compensatory mutations (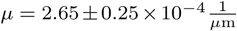, 4 nM *β*-estradiol), black markers - no rescue mutations control (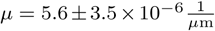, 0 nM *β*-estradiol). See Supplementary Fig. S4 for survival probability with different mutation rate. y-errors indicate Poisson distribution s.d., x-error s.d. of the mean. Inset, comparison of the ratio of compensated clones since the beginning 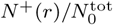 **d)** Compensated (blue line) and uncompensated (red line) clone’s average width development over time. Error bars indicate one s.d. of the mean. Width distribution for different time points of imaging on the right side, color saturation increases with time of the imaging. **e)** Efficacy of compensatory mutations, defined as the relative ratio of survival probabilities between with-rescue experiment and no-rescue control (Eq. 2). Error bars derived by Gaussian error propagation (see Supplementary Information section 0.2).

In contrast to an established null-model of a non-dense range expansion (see Methods Section for details), predicting less than 1 % presence of compensated clones, we observe an order of magnitude higher prevalence of evolutionary rescue in the experiment (Fig. 2c, inset).

Independently measuring clone widths of both uncompensated and compensated subclones, we find that the average width *w* of compensated clones (*s* = 0) grows linearly with increasing radius (Fig. 2d). In contrast, the width of uncompensated clones initially rises but then saturates for *r* ≳ 4000 and remains constant at *w*_eq_ ≈ 50 *±* 17 μm. This contrasting behaviour is also reflected in the distribution of clone widths (Fig. 2 d). While the distribution of compensated clones continuously broadens, that of the uncompensated clones remains narrowly confined around *w*_eq_.

To measure the impact of compensatory mutations on evolutionary outcomes, we compare the radiusdependent survival probability

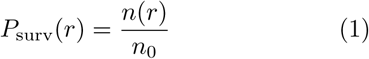

of a scenario with 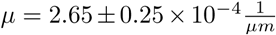 to a “no rescue” control with almost no compensatory mutations (*μ <* 10^−5^) in the time course of the experiment, with *n*(*r*) and *n*_0_ = *n*(*r* = *r*_0_) denoting the respective number of clones present at the front (see clone).

We find that the survival probability of both scenarios overlap almost perfectly for *r* ≳ 5000 (Fig. 2c). In the later phase of expansion, however, the with-rescue survival probabilities level off while those of the no-rescue control continue to decay. This divergence can be quantified by calculating the efficacy of rescue as

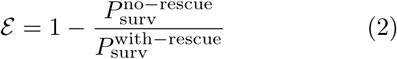

(see Fig. 2e). Here, *ε* = 0 indicates no observed difference between both scenarios while *ε* = 1 represents the limit of all surviving clones to be exclusively in the with-rescue sample. In our experiments, efficacy remains zero within errors for *r <* 4700 μm, after which it continuously rises to *ε* = 0.6 *±* 0.1 at the end of experiments.

### The impact of evolutionary rescue is delayed by an initial “window of inefficacy”

The observed delay in efficacy generates an initial “window of inefficacy” in which evolutionary trajectories seem unaffected by evolutionary rescue. Notably, its extend seems temporally decoupled from the acquisition of compensatory mutations, the majority of which having already occurred before *r* = 3000 μm (Supplementary Fig. S5). Together, this suggests an interesting consequence: The probability of treatment failure should be independent of evolutionary rescue if therapy is initiated within the window of inefficacy.

To test this prediction, we conducted a series of therapy mimicry experiments, initiating treatment either within or substantially after the window of inefficacy. In these experiments, we expanded colonies for an initial pre-treatment phase at different mutation rates for either 5 or 9 days. We then initiated treatment with Hygromycin B, halting wild-type growth, and counted resurgent growth cones after one day of post-treatment regrowth. Each cone was categorized as either compensated (+) or uncompensated (-) (see Supplementary Fig. S6). Cones that had acquired a compensatory mutation before treatment initiation but featured a remaining fraction of uncompensated cells were counted as compensated. Analogous to *P*_surv_, the respective treatment failure probabilities *P*_fail_ for each scenario is then given by

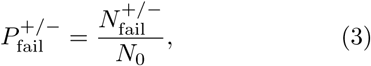

Where 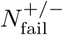 is the number of growth cones of the respective type and *N*_0_ is the number of initially inoculated clones. The total, type-independent treatment failure probability is given by 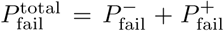. To assess the impact of evolutionary rescue on treatment failure, we can again calculate the efficacy 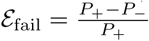.

We find that the fraction of compensated clones increases for larger mutation rates (Fig. 3a). Intriguingly, this shift in compensation status does not directly translate to a change in treatment failure probability, which remains essentially unaffected for the early treatment point (Fig. 3a). Delaying treatment until day 9, in contrast, yields a significant difference in treatment failure probabilities between control and samples with increased mutation rate (Fig. 3b). The contrast between early and late treatment time points can also be appreciated by comparing the respective efficacy values (Fig. 3c-d).

**Figure 3:**
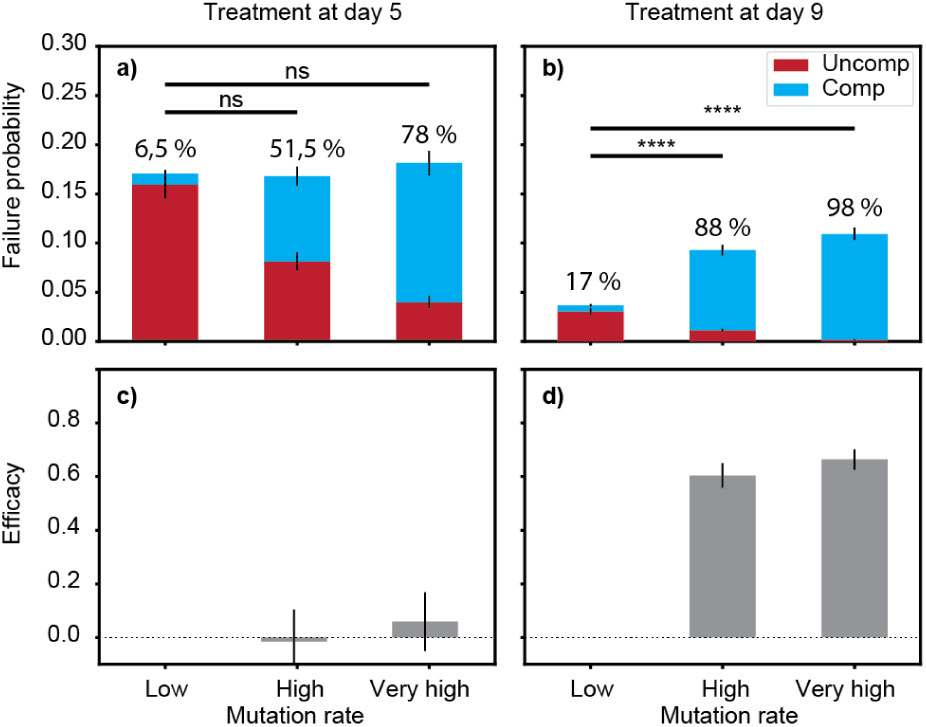
Delayed impact of compensatory mutations on treatment failure. Treatment failure probability and efficacy in experiments with different mutation rates for early (day 5 of growth) (**a**,**c**) and late (day 9) (**b**,**d**) treatment (Eq. 3). Fitness cost *s* = 0.013 ± 0.006, mutation rate low 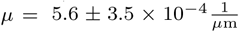, high 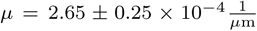 and very high 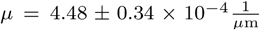. Percents represent the presence of compensated clones. Difference between the low mutation rate control and experiments with compensatory mutations are non-significant for treatment after day 5 (**b**), p-values 0.58 and 0.22 for high and very high switching rates respectively. For treatment after day 9 (**d**) respective p-values are 4 · 10^*−*22^ and 7.5 · 10^*−*30^. P-values are one-sided. Errors of *P*_fail_ indicate one s.d. of Poisson distribution, efficacy errors propagated with GEP.

### The occurrence and impact of evolutionary rescue is governed by a stable inflation-selection balance

We argue that the key phenomenon determining evolutionary dynamics, including the window of inefficacy, is the stability in width of uncompensated clones. The origin of this plateau can be rationalized by considering the interplay of the two main forces driving the change of average clonal width, i) global inflation of the population front due to radial growth and ii) natural selection. For a slower growing subclone, these forces are opposing each other with inflation increasing clone size and selection decreasing it, so that the total width change per radial expansion Δ*r* is

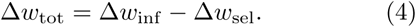

with the clonal width *w* measured as the arc length between delimiting sector boundaries (Fig. 4a,b). Inflation depends on the sector width and the current population radius as Δ*w*_inf_ (*w, r*) = *w/r* while selection is typically thought to only depend on the relative fitness difference *s* of the adjacent clones 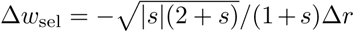. Even though there is an equilibrium clone width *w*_eq_(*r*|*s*) for any given radius at which inflation and selection forces are in balance (Δ*w*_tot_(*w*_eq_) = 0), this equilibrium is inherently unstable (Fig. 4a). However, this equilibrium can become stable if selection is width dependent, Δ*w*_sel_(*w*) (Fig. 4b). In such a stable inflation-selection balance, small clones of width *w < w*_eq_ would be inflation dominated and increase in widths while larger clones of width *w > w*_eq_ would be selection dominated and shrink (Supplementary Fig. S7).

**Figure 4:**
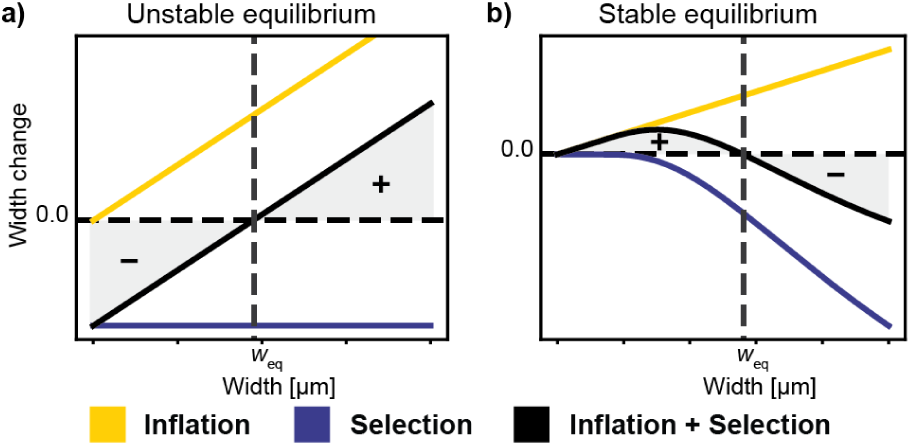
Inflation-selection balance. Schematic of unstable **a)** and stable **b)** inflation-selection balance. Yellow and blue lines represent contribution to the width change per lateral growth of purely inflation or selection. Black line is the total width change. Plus and minus indicate the widths, that will correspond to growing and shrinking clones correspondingly (see Supplementary Fig. S7 and S8).

To quantitatively test this inflation-selection balance hypothesis, we simulated both constant and width-dependent selection scenarios, modelling the trajectories of individual sector boundaries as biased 1D random walks on a radially inflating surface (Fig. 5a,b) [13, 22]. We found that a null-model of constant fitness costs failed to capture the behavior observed in experiments, even if effective selection was reduced by several orders of magnitude (see Supplementary Fig. S9). Instead, uncompensated clones exhibit the previously reported form of logarithmic spirals [20] (Supplementary Fig. S10)).

**Figure 5:**
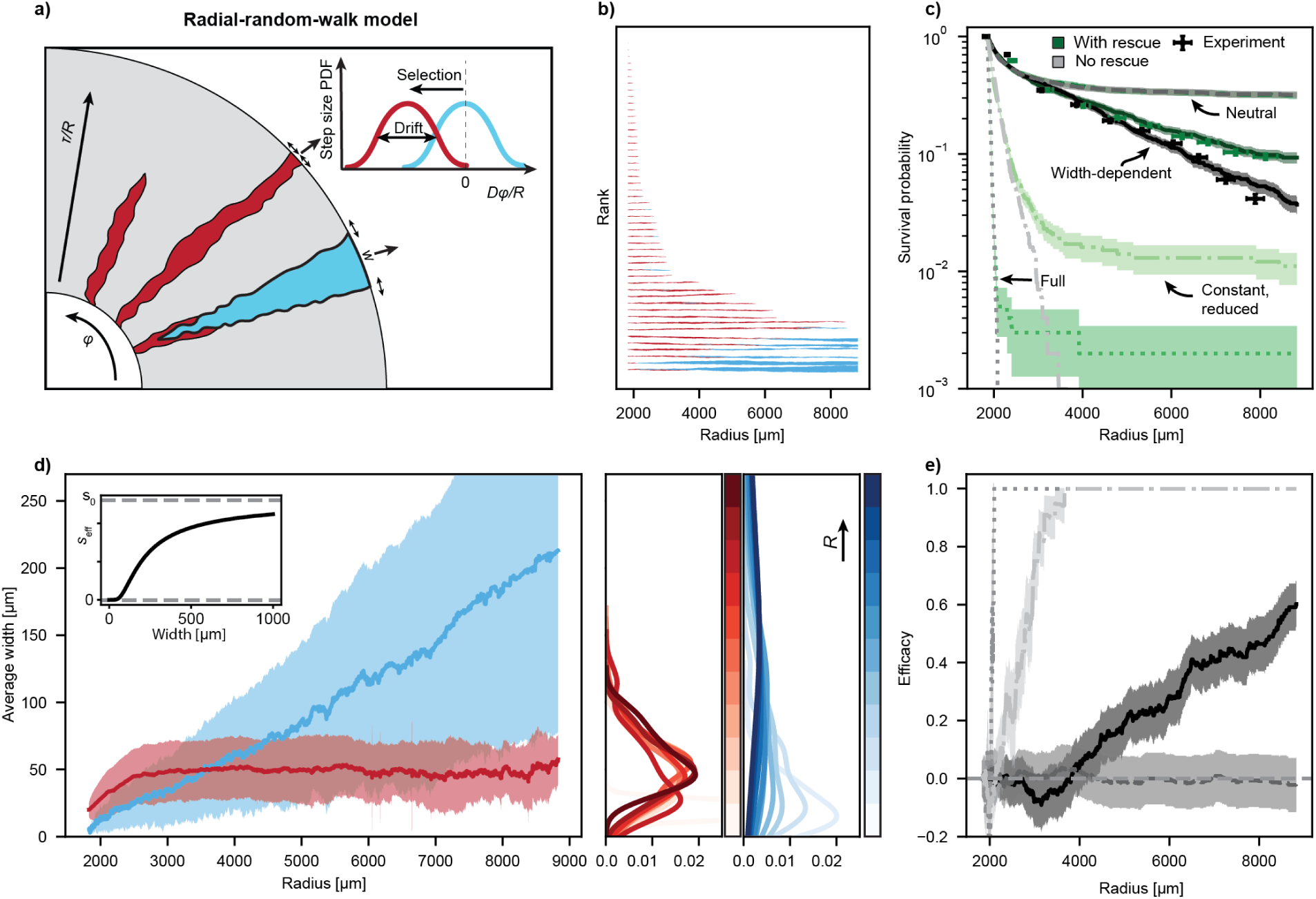
Random walk model to study Inflation-Selection balance. **a)** Schematic of null model simulation of clone boundaries as one dimensional random walk. **b)** Simulated clone trajectories. **c)** Survival probabilities for scenarios with different selection strategies (Eq. 3). Shaded area - Poisson distribution s.d. **d)** Average compensated (blue) and uncompensated clone width. Shaded area indicate one s.d of the mean. Inset, effective selection as function of clone width. Width distributions for different simulated colony radii. Color brightness increases with the radius. **e)** Efficacy of compensated clones (Eq. 2). Shaded area represent uncertainties derived with GEP.

However, using a width-dependent selection coefficient 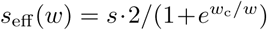 of characteristic decay width *w*_c_ (see inset in Fig. 5d) not only resulted in a plateauing and narrowly distributed width of uncompensated clones (Fig. 5d) but also captured empirical survival probabilities as well as the window of inefficacy in the efficacy of evolutionary rescue, absent in constant fitness scenarios (Fig. 5c,e). Note that the above rationale for an inflation-selection balance is robust across a wide spectrum of width dependence, solely requiring that Δ*w*_sel_(*w*) approaches zero more rapidly than inflation for declining width (see Supplementary Fig. S11). Intriguingly, we previously demonstrated that such a clone-width dependent reduction in effective selection inherently emerges as a result of collective cell dynamics in any sufficiently dense cellular populations (see Fig. 2g in reference [13]). In short, distance-dependent mechanical coupling of cell motion prevents the differential displacement required for selection to act. The exact form of this width-dependence, and with it the values of *w*_eq_, may differ substantially between systems. The concept of inflation-selection balance, however, should apply to any sufficiently dense populations. Consequently, the effects observed here are likely to extend to other types of crowded cellular assemblies, including pathogenic bacterial biofilms or solid tumors.

### Evolutionary rescue and timing dependent treatment failure in an *in silico* tumor model

To assess the relevance of our findings in the context of cancer, we conducted agent-based simulation of tumor growth using a tailored implementation of the PhysiCell platform [23]. In short, cells grow and divide in an explicitly simulated nutrient microenvironment and repulsively interact via a distance-dependent force, resulting in non-motile, overdamped motion. In our implementation, individual cells can additionally mutate at a predetermined stochastic rate and fitness effect (Fig. 6a). Cells in the population interior stop to proliferate due to a lack of nutrients, creating a peripheral growth layer.

**Figure 6:**
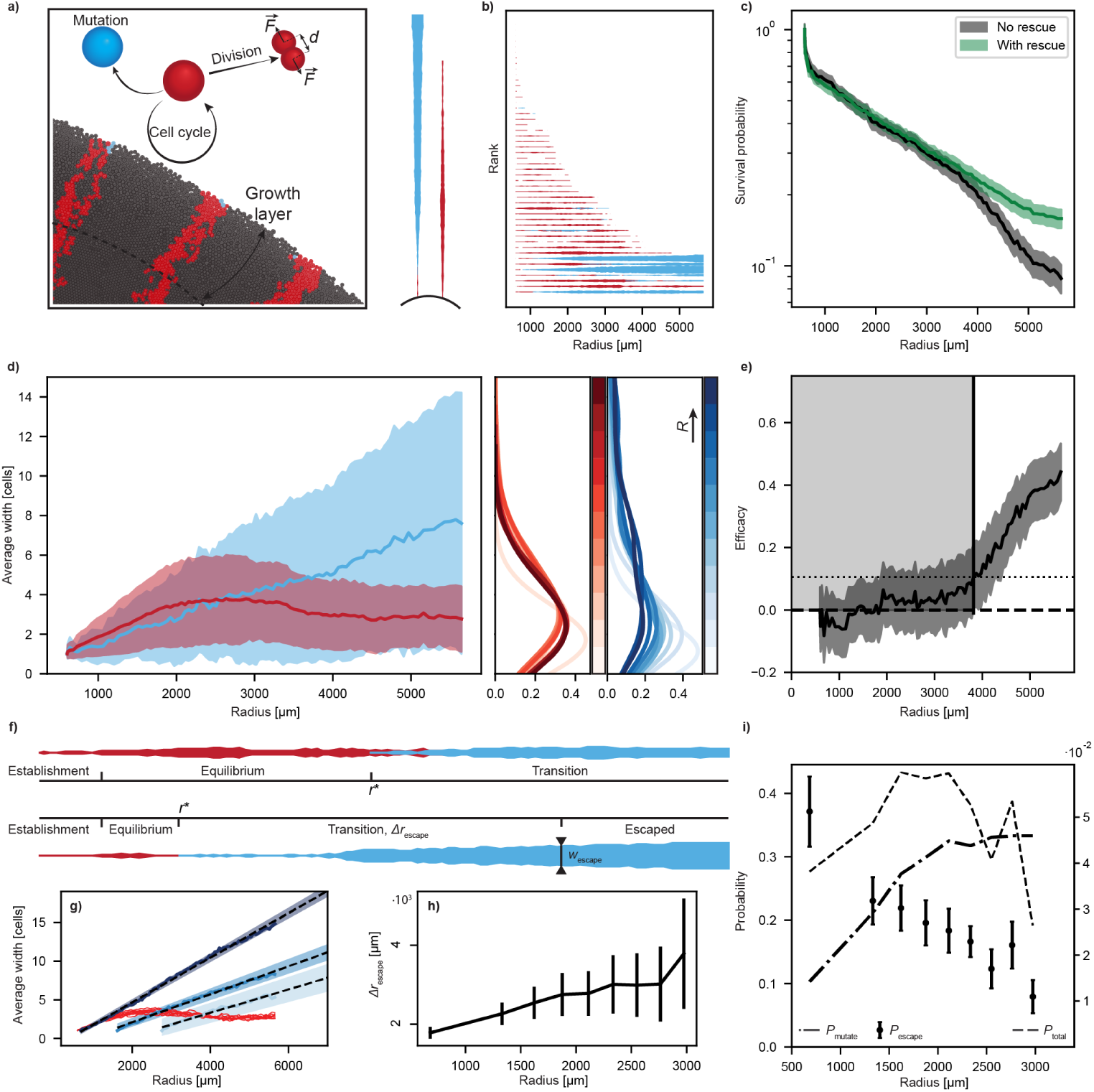
Evolutionary rescue and timing dependent treatment failure in an *in silico* tumor model. **a)** Schematics of the simulation set-up. **b)** Simulated clone trajectories. **c)** Survival probability as the function of colony radius (Eq. 3). Shaded area indicate one s.d. of Poisson distribution. **d)** Average compensated (blue) and uncompensated (red) clone width in cell diameters and its distribution as function of colony radius. Shaded area indicate one s.d. of the mean. **e)** Compensatory mutation efficacy (Eq. 2). Shaded area indicate errors propagated by GEP. **f)** Simulated clone trajectories with identified phases. **g)** Solid lines - average clone width development with colony growth for different mutation radii. Dashed lines - linear fit to the width of compensated clone size. Shaded areas represent uncertainties in the fitted lines. **h)** Δ*r*_*esc*_ as the function of mutation radius. Error bars derived from the uncertainties of linear fit to average width. **i)** Filled circles - probability to grow above escape width *w*_*esc*_ = 6 cells in 2500 m of colony growth after mutation (error bars indicate one s.d. of Poisson distribution), dash-dotted line - probability to get a mutation (with mutation rate *μ* = 0.1), dashed line - probability for a clone to get a mutation and eventually grow to the size that makes its probability to survive higher than 99 %. X-axis mutation radius.

Similar to our experimental assay, we simulated the expansion of 2D *in silico* tumors starting from a mixed inoculation of resistant but slower-growing cells interspersed at a low fraction (16.6%) into a background of faster growing wild-type cells (Fig. 6a).

Resulting *in silico* tumor populations exhibited a striking resemblance with our experimental yeast model system (Fig. 6b), featuring the same thin yet persistent streaks of uncompensated clones rescued by neutrally expanding compensated subclones. Following an analysis pipeline equivalent to that applied to experimental data, we find that simulations capture the full finger print of inflation-selection balance, including the experimentally observed boost in resistant subclone longevity and the resulting increase in treatment failure probability (Fig. 6c-e).

With the ability to track the composition, width and eventual fate of each individual clone with singlecell precision on an intra-generational time scale these agent-based simulations are ideally suited to quantitatively test our hypothesis of a finite equilibrium clone width of uncompensated subclones as an emergent phenomenon.

We find that the life history of a clone can be divided into four characteristic phases (Fig. 2f). At the beginning of the initial *establishment phase*, uncompensated clones are very small by definition, similar to a *de novo* resistant clone just after acquiring the resistance-conveying mutation. During this phase, the probability of a clone to go extinct due to random genetic drift is high.

Those clones not succumbing to genetic drift right away enter the proposed *equilibrium phase*, in which the width of a clone is held constant by the opposing forces of natural selection and peripheral inflation. In our simulations, we find this finite equilibrium width to be 39.24 *±* 22.32 μm, or *w*_eq_ = 3.27 *±* 1.86 cell diameters. While clones in this phase can still fluctuate to extinction, the rate is drastically reduced in comparison to the initial *establishment phase* due to an increased average clone size. This stabilization at a small but finite clone width can now be interpreted as the root cause for the narrow yet persistent streaks of uncompensated clones observed Fig. 1g.

The acquisition of a compensatory mutation then initiates a *transition phase*. As before for the uncompensated clones, *de novo* compensated subclones are very small and likely to be driven to extinction by genetic drift. However, being neutral with respect to the wild-type and even having a fitness advantage over their neighboring uncompensated ancestor, compensated clones are constantly driven towards larger widths by inflation.

Once a clone becomes larger than 6 cells, with width *w > w*_escape_ *≈* 72 m, probability of genetic drift overcoming inflation becomes smaller than 1%. Clones reaching this *escaped phase* not only persist indefinitely but continue to grow in lateral width until treatment is initiated at an arbitrary point in the future.

A direct consequence of the observed quasi-stability of slower-growing clones is that compensatory mutations can occur over a wide range of population radii. This raises the question of how the probability of a compensated clone to permanently escape selection, and ensuing long-term persistence, depends on the point *r* = *r*^*^ at which the compensatory mutation occurred.

Our agent-based simulations allow us to trigger compensatory mutations at a predefined radius *r*^*^ and then measure the resulting escape probability *P*_esc_(*r*^*^) (Fig. 6b,d).

We find that *P*_esc_ is highest for early mutations and then declines with increasing *r*^*^. The origin of this dependence can be understood by considering the expected expansion 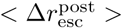 it takes a *de novo* compensated mutant to grow to escape width (Fig. 6g,h). The slope of the average width of radially expanding uncompensated clones is inversely correlated to *r*^*^, due to a continuous decrease in front curvature. As a result, compensated clones with greater *r*^*^ spend more time in the transition phase during which they can still be driven to extinction by random genetic drift.

Combining the evolutionary dynamics of both uncompensated and compensated clones, we can now calculate the total probability that a resistant subclone survives until and then escapes due to a compensatory mutation at *r* = *r*^*^ (Fig. 6i). Because of the initial inflation-dominated part of the inflation-selection balance, mutation probability is increasing initially and then saturates. We find that the opposing behaviors of an increasing mutation probability and a declining escape probability result in a peak of combined rescue probability. This peak could also exist in the inflation-dominated part of the constant selection scenario. But as it inherently produces an unstable equilibrium, and clones smaller than equilibrium width will shrink, inflation has to be very strong for this scenario to be achieved for *de novo* mutants. For example, with a fitness cost of 1% the colony radius would have to be as small as only three cell diameters for a single-cell clone to be in the inflation-dominated regime. On the other hand, in comparison to a predicted by the null model decay of the mutation probability, its saturation widens the *P*_tot_(*r*^*^). As a result, it also increases cumulative rescue probability, corroborating our notion of enhanced drug resistance evolution in dense populations.

## Discussion

In this work, we introduce a new experimental approach, based on a tailored microbial model system featuring fluorescence-coupled synthetic mutations, to systematically study evolutionary rescue dynamics in dense cellular populations and resulting consequences for persistent drug resistance.

Tracking the complete evolutionary trajectories of individual clones, we identify a novel inflation-selection balance, in which the counteracting effects of peripheral population inflation and a clone-width-dependent selection pressure result in a quasi-stable equilibrium width of slower-growing resistant clones. As a result, these lineages persist at the expanding population front, improving their chance to acquire a subsequent cost-compensatory mutation.

Intriguingly, inflation-selection balance is a double-edged sword. On the one hand, it increases the *prevalence* of evolutionary rescue, while, on the other, it creates an initial “window of inefficacy” during which the *impact* of evolutionary rescue on the probability of treatment failure remains low.

Using an agent-based *in silico* model of tumor growth, we additionally demonstrate that inflation-selection balance only requires a minimal set of ingredients, also present in solid tumors. We conclude our study by leveraging our simulation platform approach to investigate how the timing of compensatory mutations impacts long-term treatment success.

The minimal approach presented here has several important limitations that need to be considered when extrapolating our findings to other types of dense populations. While our well-controlled yeast model system is ideally suited to investigate the fundamental effects of density-driven growth on the evolutionary dynamics of dense populations, it does not capture many of the intricacies present in other settings, especially those of solid tumors. This suggests future avenues of research, studying how more complex cellular features, such as active migration, cell-cell adhesion or heterogeneous mechanical properties modulate the fundamental processes discussed in this work.

It should be noted that our study focuses on the evolutionary rescue of a pre-existing resistance mutation. Alternative trajectories to two-step resistance, such as the hitchhiking of a resistance mutation on an independent driver mutation, have not been included [24]. However, our experimental strategy and the presented computational framework could be generalized to investigate these alternative scenarios.

Notably, our observations might not be exclusive to dense populations. While the necessary ingredients for inflation-selection balance inherently emerge from density-driven growth, they might also be generated by other forms of negative size-dependent selection, such as it has been described for mutualistic scenarios [25]. We predict the existence of an inflation-selection balance to be robust across a wide range of system parameters, including the shape of width-dependence of selection, and systems.

Of particular interest will be to extend our investigations to three-dimensional range expansions, such as matrix-embedded microbial spheroids and cancer cell tumoroids.

In conclusion, our work is a crucial step towards understanding the intricate evolutionary dynamics in dense cellular populations as an emergent phenomenon in actively proliferating granular matter. This study is, to our knowledge, the first to combine the effects of two of the most fundamental features of these systems, growth-driven radial expansion and collective dynamics, to investigate their impact on critical evolutionary processes, such as evolutionary rescue and drug resistance evolution.

Going forward, the uncovered interplay between inflation-selection balance and multi-step adaptation should be considered when predicting critical consequences of cellular evolution, such as drug resistance. Finally, the insights gained may serve as an important stepping stone towards new evolution-based treatment strategies [4, 26–28].

## Methods

### Strains

All experiments in this work were conducted with the non-motile yeast *S. cerevisiae*. The used strains, yJK26 (resistant “mutant” with synthetic mutation system) and yMG10 (“wild-type” background), were constructed on the W303 laboratory strain and are based on the common ancestor yJK19, expressing the β-estradiol inducible pSCW11-Cre-EBD recombinase construct from pDL12[29].

yJK26 and yMG10 feature the same synthetic mutation cassette *P*_ENO2_*-loxP-ymCherry-kanMX-loxP-FP2-UBQ-cyh2r*, only differing by the secondary fluorescent protein *FP2* behind the cassette, which for yJK26 is *ymCerulean* and for yMG10 *ymCitrine* (see following section for details cloning of the synthetic mutation system). In both cases, the fluorescent protein is coupled via a proteolytically cleavable ubiquitin linker to *cyh2r* (*CYH2Q37E*), conveying resistance to the translational inhibitor Cycloheximide. To construct yJK26, we first introduced the synthetic mutation cassette (yJK20) and subsequently replaced the Nourseothricin resistance marker, originally introduced into yJK19 as selection marker for *cre-EBD* insertion, with a HygR resistance from pAG32 (available from www.addgene.com, #35122). Consequently, yJK26 (and its converted version yJK26c) are constructively resistant against Hygromycin B, while the converted version of yMG10 (yMG10c) is used as Hygromycin-susceptible “wild-type” strain in our experiments. All insertions were achieved by amplifying the insert via standard PCR, followed by Lithium Acetate transformation and selection. Correct insertion was verified by cross-junctional colony PCR of positively selected clones. The genotypes of the strains are specified in the Supplementary Table 1.

### Construction of synthetic mutation cassette

The synthetic mutation cassette was constructed on the basis of pMEW90 (a kind gift of the lab of Andrew Murray, Harvard)[30]. Initially, all parts were amplified via PCR and assembled via Gibson cloning to yield pMG8. To obtain pJK19, we replaced the *ymCitrine* in pMG8 with *ymCerulean* via Gibson assembly (for plasmid maps see Supplementary Fig. S12 and S13).

### Evolutionary rescue assay

Evolutionary rescue experiments were conducted subjecting yJK26 and yMG10c to spatial competition on 2 % agar YPD plates. From the colonies grown from a single cell on the agar plate for 2 days parts of the single colonies were picked and cultured in liquid YPD medium overnight. The next day cells were reinoculated into fresh medium and regrown for about 3 hours. The two strains were mixed in the ratio 1:9 resistant to susceptible, estimating cell concentration by *OD*_600_ measurement. 1 μL of mixed cultures of *OD*_600_ ≈ 20 were then inoculated on the 6-well plates filled with 15 mL medium and air dried. Image of the colony ∼ 1 hr after inoculation shown in Supplementary Fig. S14. Cycloheximide and *β*-estradiol were premixed in the medium in desired concentrations (50 nM for CHX, and 2, 4 or 6 nM for BED). Colonies were grown for 5 or 9 days prior to treatment and imaged on a daily basis. At the treatment day Hygromycin B was applied by pipetting 90 μL of 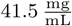 stock as small drops at the edges of the wells. Total of 92 colonies were grown, with at least 5 colonies in the same chemical environment. Treatment after 5 days was initiated for 6 colonies of each condition (50 nM CHX + 0, 4 and 6 nM BED and 0 nM CHX + 0 nM BED (compensated inoculum)), except 0 nM CHX + 0 nM BED (uncompensated inoculum), where only 5 colonies were treated. Treatment at day 9 was done for 18 colonies with 50 nM CHX + 0 and 4 nM BED, 17 colonies with 50 nM CHX + 6 nM BED, 6 and 5 colonies with no chemicals for uncompensated and compensated inocula correspondingly. Initial number of clones is estimated by manually counting clones from the single-cell resolution images. From 12 colonies we measure mean of 156 clones per colony, that results in the total of ≈ 14500-15000 clones.

Fitness differences between the strains under different CHX concentrations were measured using the final images of colonies grown from the inoculations with low fractions (2.5-10%) of faster-growing cells (yMG10c) in the slower growing (yJK26). Opening angle of the faster growing clones were measured at different radii of the colony and fitted with Eq. 10 from the reference [20] to extract growth rate difference. For 50 nM CHX fitness difference between yJK26 and yJK10c is Δ*s* = 1.26 ± 0.64% (measured using 14 non-interfering clones). For yJK26 and yMG10c under no CHX Δ*s* = 0.09 ± 0.08% (measured using 8 non-interfering clones). Fitness difference between yJK26c and yMG10c is within the errors of the method zero, and both strains are considered to have the same fitness. Colony examples and measured values are presented in the Supplementary Fig. S2 a,b.

Mutation rates of yJK26 were estimated by measuring frequency change of mutated clones throughout colony growth. 1 μL of yJK26 cells in YPD with *OD*_600_ ≈ 20 were inoculated onto the agar plates containing different concentrations of BED and imaged daily. Frequency of compensated mutant at the colony periphery was measured and plotted over colony’s radius, averaged over 6 colonies (see Supplementary Fig. S15) for 0, 2 and 4 nM BED and 12 colonies for 6 nM BED. Considering both clones neutral, change in the frequency only occurs due to mutations and can be described by the rate equations. This results in the change in frequency of blue clones

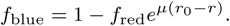

Where *f*_red_ - frequency of red clones, *μ* - mutation rate, *r*_0_ - initial radius, r - current colony radius. Fitting the function above to the frequency change, mutation rate per cell per radial colony growth was extracted. Orthogonal distance regression was used for fitting and calculating uncertainties of the mutation rate. See Supplementary Fig. S2 c,d.

### Imaging and analysis

Colonies were imaged using Zeiss Axiozoom V16 fluorescence microscope with PlanNeoFluar Z 2.3x/0.57 objective for imaging right after the inoculation and after 1 day of growth and the PlanApo Z 0.5x/0.125 for all other time points. A custom semi-automated pipeline was used to process the images. First, images were segmented with a Machine Learning based platform Ilastik [31]. The algorithm was trained on randomly picked images from different days (training data is available upon request). Colony microscopy images with segmented outputs next to them are shown in the Supplementary Fig. S16. Segmented images were further analyzed via a custom MATLAB algorithm (code available on gitLab).

### Clone trajectories

Clone trajectories (Fig. 2b) are reconstructed by assigning labels to the each individual clone at the periphery. For labeling, angular positions throughout different days of the imaging were used. Image after one day of growth was taken as the reference. For each next day clone angular positions were compared to clone positions one day before to assign IDs. Full trajectory tracking was only done for illustrative purposes for one colony and was not needed in the quantitative analysis presented in the paper. Nearest neighbor method used here to assign clone IDs only works robustly for very round colonies, and would need further improvement to be applied for all of the grown colonies.

### Description of the random walker model

We treat the boundaries of each resistant clone as independent non-interacting one dimensional random walkers [19, 32]. Each individual walker is a solution to the diffusion equation. Diffusion coefficient is the measure of genetic drift in the system that is represented by the random step that follows Gaussian distribution with variance [19]:

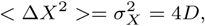

and the step:

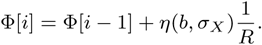

Where *η*(*μ, σ*) is normally distributed noise with mean *μ* and variance *σ. b* is the bias of the walk, which is zero for the neutral clones, and can be positive or negative number for clones with a fitness deficit, depending on the side of the sector that the random walker is representing. To simulate clonal extinction we conditionalize neighboring random walkers to be annihilated when they cross. Bias of the random walk *b* is calculated using equal-time argument [19, 20]:

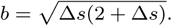

To support our argument of inflation-selection balance emergence from collective cell dynamics we use effective width-dependent selection coefficient. Thus *s* in the bias of the random walk is *s* = *s*(*w*). Exact analytical form of the effective selection depends strongly on the amount of pushing in the colony, that might be changing in time and is too complex to derive. We argue that the exact shape is not important for balance to emerge (see Supplementary Fig. S11). To guide the shape of *s*_eff_ for use in the simulation we use experimental data from Fig. 2g from the ref. [13]. *s*_eff_ used in the null model has the form

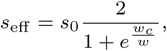

where *w*_*c*_ is the critical width analogous to the critical width in ref. [13] and is a free parameter.

### Parameterization of the random walker model

To parameterize the random walk model, we use measured initial radius of the colony after inoculation and measured fitness cost *s*_0_ (see Supplementary Information section 0.3). To obtain the value for *w*_*c*_ in effective selection we compared the equilibrium width of slower growing clones to the experimentally measured one. For the fitness cost of *s*_0_ = 0.013 critical width of 280 μm matched experimental data. To estimate genetic drift in the system, we tuned diffusion coefficient together with initial clone size distribution to match the experimental clone survival probability (Supplementary Fig. S17, S18. Diffusion coefficient to match experiments is 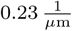. Strength of the drift controls the shift of survival probability, and initial width distribution - how steep is the initial drop of the probability, when most of the clones are of very small size. We end up with initial size distribution that follows Gaussian distribution with mean of 20 μm and standard deviation of 5 μm. For comparison mean width of red clones after 1 day is 26 ± 12 m. Larger then experimentally observed initial clone size needed to match the drop of survival probability in the null model can be caused by smaller cell density in the experiment right after inoculation and clone widening during the initial exponential growth phase and filling the gaps in the center of the colony (see Supplementary Fig. S14). Analogously we also compare the results with different mutation rates (see Supplementary Fig. S19). Mutation rate used in the simulations 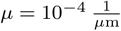 is in a good agreement with experimentally measured value of 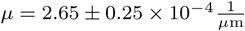.

### Agent-based simulations

To show that only minimal ingredients are needed for the results, we obtained in the experiments, agent-based model (ABM) is used to grow virtual dense populations. We grow our virtual colonies using a modified version of the open-source platform for 2D and 3D tumor growth PhysiCell (Version 1.8.0) [23] with BioFVM [33] to solve the transport equation.

It simulates cell growth, including mechanical cell-cell interactions, on top of the chemical microenvironment that is modeled via diffusion of chemicals in the system. Cell division and physical forces exerted on the neighbors lead to densely packed population. Agents continuously consume nutrients diffusing to the system from the outer edge with circular Dirichlet boundary conditions. It results in the emergence of the growth layer, when population density reaches certain threshold and nutrients in the center are depleted (see Supplementary Video 2). These features, which we speculate to be essential to the dynamics we observe experimentally, thus emerge inevitably in this system which includes very few fundamental ingredients.

Each cell in the simulation carries its own characteristics, defined by its mechanical and chemical environment. These characteristics include cell size, cycle progression, growth rate and mechanics. We initialize the system similar to the initial inoculations in experiments, having a hollow dense ring of cells, with sufficiently spaced single slower-growing mutants at the periphery. Growth of both cell types depends on the nutrient concentration, slowing down gradually until the growth is completely halted for very low concentrations. For parameterization of the simulation see Supplementary Information section 0.3.1.

Ratio of cell size to the colony radius is different from the experiment due to computational limits. Virtual populations are grown up to ∼ 2 · 10^6^ cells, that is tenfold radial increase from the starting colony.

